## Supplementary material for "Amalgam plays a dual role in controlling the number of leg muscle progenitors and regulating their interactions with developing tendon": Full revision answers

**Manuscript number:** RC-2023-02319
**Corresponding author(s):** Cedric, Soler

#### 1. General Statements [optional]

*This section is optional. Insert here any general statements you wish to make about the goal of the study or about the reviews.*

First, we would like to extend our gratitude to all reviewers for their supportive and enthusiastic feedback, which acknowledges our study as an interesting, well-executed, and well-documented contribution to the field. We are also pleased that the novelty and significance of our work have been recognized and appreciated.

As highlighted by reviewers 2, 3, and 4, our research represents a substantial advancement in understanding the mechanisms that coordinate the development of different cell types. Our findings have broader implications for developmental biology. We also thank the reviewers for their valuable insights, which have significantly improved the overall readability of our manuscript. We have carefully considered all minor corrections and text modifications they had suggested and made amendments accordingly.

The reviewers proposed several complementary experiments to enhance and clarify our points. We have conducted most of these experiments, with one exception (detailed below), and incorporated the corresponding results into this revised version of the manuscript.

Additionally, reviewers agreed that higher resolution images depicting the interactions between tendon and myoblast membranes would strengthen our manuscript. In response, we are pleased to present new high-resolution images of Ama::EGFP localization with respect to tendon and muscle cells, obtained using Zeiss Airyscan technology. We also provide new images using newly generated flies that allow simultaneous observation of both myoblast and tendon membranes.

We believe these modifications substantially enhance the quality and interest of our results, as already highlighted by the reviewers.

#### 2. Point-by-point description of the revisions

*This section is mandatory. Please insert a point-by-point reply describing the revisions that were <u>already carried out and included</u> in the transferred manuscript.*

****Referees cross-commenting**** :

***All reviewer agree*** *that "this is an interesting study that is well done and well documented. I agree with reviewer 1 that the study would further benefit from better imaging of the cellular extensions of tendons and myoblasts to see how both cell types interact."*

**Reply**: We agree with this point. To address it, we analyzed leg discs from Sr-Gal4>UAS-myrGFP line (labelling tendon membranes) crossed with a newly generated line R32D05::CD4TdTomato (myoblast specific expression of membrane tagged Tomato protein). Using confocal Zeiss Airy Scan technology, we generated high resolution images for which both tendon cell extensions and myoblast membranes are simultaneously visualized. These images are included in the new Figure 2 (O, P and O', P'). To be noticed: as we also provide new high resolution images of Ama::EGFP and Nrt localizations (Fig. 2 M, N and M', N'), we removed images of zoom in of 5h APF leg disc.

***REVIEWER 1:***

*Moucaud et al carried out single cell sequencing on myoblasts from the developing drosophila leg muscles, focusing on gene expressions overlapping with tendon and muscle cells. This study proposes that neuronal cell adhesion molecules Ama and Nrt interact in myoblast and tendon adhesion to support tendon and in proliferation of muscle progenitors. This study traces Ama and Nrt expression with various drosophila mutant strains and provides evidence to support its claims using single cell sequencing, immuno-fluoresence and in situ hybridisation.*
*The authors report novel markers to study the interactions between muscle and tendon progenitors in the Drosophila leg provide convincing evidence of their functions in muscle and muscle and tendon formation.*
**Reply:** We are grateful to the Reviewer 1 for his/her supportive comments on the quality of our work.

***Point-by-point reply:***

- ***Page 2 "cell types****..." might be worth including other cell types such as vascular/endothelial if listing all cell types in the limb, as the sentence is suggesting.*
**Reply**: "blood vessels" have been added as components of the limb musculoskeletal system.

- ***Reviewer's comment:*** *The authors discuss the interactions between the myoblasts and tendon cells but do not show any cellular resolution of the interaction between the cells and the secreted adhesion proteins. It would enhance the manuscript if the authors could show high resolution images of these cellular interactions with the secreted protein in vivo.*
**Reply**: see reply to referees cross-commenting about newly generated high-resolution images shown in Fig. 2.

**Minor typographical :**
*- Lots of examples of definite article (the) missing throughout the text.*
**Reply**: The text has been edited and missing articles added

*- Second line of Abstract does not flow: Ama encodes secreted proteins to "Ama encodes a secreted protein"* the correction has been made accordingly

*- 2nd para Intro- this para is essentially discussing vertebrate limb muscle/tendon precursors, although includes a non-vertebrate citation. It could be helpful to (briefly) compare/contrast the non-vertebrate vs vertebrate literature on this topic.*
**Reply**: Indeed, this paragraph is primarily focused on the development of the musculoskeletal system in vertebrates. The comparison (from a molecular standpoint) with the muscle system of the Drosophila leg appears in the following paragraph. For clarity, we have included a brief, more general description (end of second paragraph) about the muscle/tendon system in

Drosophila to highlight certain divergences between vertebrate and invertebrate systems and to introduce the subsequent paragraph.
- "*in limb of chick embryo add "the limb"* the correction has been made accordingly
- *p6 because these two antibodies were raised in rabbit, as the Twist antibody, needs some additional explanatory text.*
**Reply**: We have modified the text to give a more accurate explanation: "Because these two antibodies were raised in rabbit, as was the Twist antibody, we could not use this latter to visualize the myoblasts"
- *P9 discussion creeping into results section-with some speculation on Ama forming homophilic adhesions which has not been experimentally tested.*
**Reply:** Because we chose to submit this work as a short format paper, Results and Discussion sections are indeed combined. However, we agree that homophilic adhesion properties of Ama have been shown only in cell culture and not tested in physiological context. To clarify this point, we have modified the corresponding part of the text and only suggest that Ama could directly bind to membranes through its putative GPI modification as proposed by Seeger et al.

**Reviewer #1 (Significance (Required)):**
*The authors report novel markers to study the interactions between muscle and tendon progenitors in the Drosophila leg provide convincing evidence of their functions in muscle and muscle and tendon formation.*

*- Ama depletion affects both viability and the proliferation rate of leg disc myoblasts (in a Nrt-independent way) Does it have similar role in tendon precursors? Could the authors provide any evidence of apoptosis given proposed role of Ama in glial cells?*
**Reply:** As asked by Reviewer 1, we have tested these two points and included the results in suppl figure 3 (A-C). As expected, the proliferation rate of tendon cells is not affected as we have previously showed that tendon cells are post-mitotic cells (Laurichesse et al. 2021). Moreover, we now show that Ama depletion does not lead to apoptosis of tendon cells. See Supp. Figure 3 (A-C), the main text has also been modified accordingly.

***REVIEWER 2:***

*In this well-written, comprehensive, and interesting manuscript, the authors study the molecular circuitry that supports the coordinated activity of tendon cells and myoblasts during development. As the authors themselves point out in the introduction, the assembly of tissues within the musculoskeletal system provides a particularly attractive system in which to study how different cell types coordinate their behaviours to form higher-order structures. Using single-cell transcriptomics, the authors first identify the cell adhesion molecule Ama and transmembrane protein Nrt as enriched in Drosophila myoblasts and tendon cells. Their transcriptomic data suggest that Nrt is specifically expressed in the tendon cells while Ama is expressed in both. They support these data with a variety of in situ, antibody, and endogenous stainings. Using a series of genetic manipulations, they then convincingly show that Ama controls the total number of myoblasts during the larval stages: Ama knockdown is associated with both decreased proliferation and increased apoptosis of myoblasts. Ama's role in regulating myoblast number is shown to be independent of Nrt and likely under the control of the FGF/RTK pathway. Finally, the authors show that the loss of either Nrt or Ama activity is associated with a loss of adhesion between myoblasts and tendon cells and with the stunted growth of the long tendon. Thus, their data point to Ama playing dual roles in muscle development by regulating both myoblast number and cell adhesion.*

*I very much enjoyed reading the paper, which I think makes an important contribution to our understanding of both the developing musculature and inter-cell-type coordination during development more broadly. I have only a handful of grammatical errors to point out.*
**Reply:** We appreciate these enthusiastic and supportive comments, and we would like to thank the Reviewer for highlighting the broader contribution of our work to the understanding of the mechanisms of coordination between different cell types.

***Point-by-point reply :***
*- Grammar: 'This prompted us to use Drosophila model to search' should read 'This prompted us to use the Drosophila model to search'*
*- Grammar: '...identify Neurotactin (Nrt) and its binding partner, Amalgam (Ama) as candidates...' should read '...identify Neurotactin (Nrt) and its binding partner, Amalgam (Ama), as candidates...'*
*- Grammar: 'As tendon precursors in leg disc' should read 'As tendon precursors in the leg disc'.*
*- Grammar : '...we performed a series of in situ hybridization...' should read '...we performed a series of in situ hybridizations...'*
*- Grammar: 'Because these two antibodies were raised in rabbit, as the Twist antibody' should read 'Because these two antibodies were raised in rabbit, as was the Twist antibody'*
*- Grammar: 'Statistical analysis reveals an increase myoblast total number when overexpressing an activated ERK' should read 'Statistical analysis reveals an increase in the total number of myoblasts when overexpressing an activated ERK'*
*- Grammar: The following section header needs rephrasing: 'Ama, potential downstream effector of FGF pathway in the regulation of myoblast number'. Maybe 'Ama is a potential downstream effector of the FGF pathway in the regulation of myoblast number' or Ama: a potential downstream effector of the FGF pathway in the regulation of myoblast number.*
*- Grammar: 'whereas its reduction (UAS-StyRNAi) lead to more myoblasts' should read 'whereas its reduction (UAS-StyRNAi) leads to more myoblasts'*
*- For clarity 'The expression of the constitutively active form of ERK could rescue the phenotype of Ama depletion in glial cells (Ariss et al. 2020)' might read better as 'Previous work has shown that the expression of the constitutively active form of ERK can rescue the phenotype of Ama depletion in glial cells (Ariss et al. 2020). Therefore, we tried...'*
*- Grammar: 'showed a significant higher number of myoblasts compared' should read 'showed a significantly higher number of myoblasts compared'*
*- Grammar: 'Another, not exclusive, possibility' should read Another, non-mutually exclusive, possibility'.*
*- Grammar: 'We measured the length of the tilt relatively to the length' should read '. We measured the length of the tilt relative to the length'*

**Reply:** All the modifications suggested above by Reviewer 2 are now included in the text.

***REVIEWER 3:***

*Myoblast and tendon precursors stem from different developmental origins. Hence, they need to find each other to build a functional muscle-skeleton. How they do so is an exciting biological problem, not only for this reviewer who is working on Drosophila muscle development, too.*

*As we currently understand little about how myoblasts communicate with tendons during development, I find this manuscript a generally interesting contribution unravelling a new mechanism of cell-cell communication between these two cell types. It proposes a role for 2 interesting proteins that are little studied. Furthermore, Drosophila leg muscle-tendon development is complex and results in an intricate final architecture. Thus, a better understanding of its molecular mechanisms of development is exciting to this reviewer and to the muscle and tendon fields.*

**Reply:** We express our gratitude to Reviewer 3 for his/her keen interest in our work and for emphasizing its significance within the field of developmental biology.

***Point-by-point reply :***

***-** While some Ama mRNA expression in myoblasts was confirmed with in situ hybridisation, it was also shown that Ama mRNA is expressed in other sources including tendon precursors. As the interesting AmaGFP protein overlapping with the developing tendon cells is found at some distance from the myoblasts, the source for this Ama protein population is not entirely clear. To identify if it is secreted from myoblasts I suggest to stain for Ama-GFP in the muscle-specific Ama knock-down discs at 5h APF. This could use the late knock-down condition.*

**Reply:** In Suppl Fig.2 in a close-up view of the femur region, we show that at 5h APF Ama is transcribed in addition to myoblasts also in the developing tilt tendon. This tendon associated Ama expression is specific as it is detected after myoblast specific Ama knockdown. Thus, at 5h APF, the Ama-GFP signal detected at the interface of muscle and tendon precursors could in part correspond to Ama secreted by the tilt tendon cells. However, we also observed clear Ama-GFP signal at the interface of myoblasts and tendon precursors at 0h APF when Ama is not yet transcriptionally activated in tilt tendon precursors (not shown). Thus, we are confident that the myoblasts are the main source of secreted Ama protein that ensure close proximity of myoblast and tendon precursor cells. A view supported by the loss of myoblast-tendon cell proximity in myoblast-specific Ama knockdown. However, to clarify this point, we immunostained myoblast-specific Ama knock-down discs for the Ama protein in at 5h APF as suggested. As stated in the text, we were concerned that GFP tag could influence the life-time of the Ama protein, as GFP itself is pretty stable. This is why we used anti-Ama antibody kindly provided by Dr. Silman to determine whether myoblast-specific Ama knockdown (using R32D05-Gal4 driver) would completely abolish Ama protein at 5h APF. We indeed observed a strong reduction of Ama protein at this stage indicating that the contribution of Ama protein from tendon cells is minimal (but cannot be completely excluded), with myoblasts remaining the major source at this stage. This new result is now presented in Suppl Fig. 2M-P. This result is also in accordance with our new result showing that tendon-specific AmaKD has no effect on tendon growth (see reply to the comment below regarding tendon length). In light of this new result, we have modified the text accordingly in the corresponding paragraph (p5-6).

*- Generally, it might be useful to move the part of Figure 2 that shows the Ama-GFP Nrt co-staining to the later part in the text that addresses the interaction of both cell types and keep the autonomous Ama role in muscle for the start of paper only.*

**Reply:** We have indeed debated extensively about this possibility before submitting this work. While such a presentation would have some logical coherence, it also has the disadvantage of having to resume, at least partially, the expression of Ama, leading to certain redundancies. Additionally, we chose to begin with a comparison of the new myoblast transcriptomic data with pre-established tendon data to highlight the presence of ligand-receptor pairs. In this context, it seemed to us more pertinent to present the expression patterns of Ama and Nrt together in the initial figures.

*- To quantify the interaction of the myoblast cell membranes and the tendon cells better it would be useful to combine sr>CAAXmCherry with a myoblast membrane maker (possibly Him-CD8-GFP or use R15B03-Gal4 with R79D08-lexA).*

*This could also improve the "mean distance" measurements. As currently presented, it is not so clear how the mean distance was measured. It could be helpful to indicate some examples in zoom-in vies on Figure 5. Does a distance of 4 µm in wild type mean that the myoblast is not touching the tendon precursors, or is only the myoblast nucleus that is Twi positive at this distance?*

**Reply:** We are grateful to this reviewer for its relevant suggestion. Thus, as stated above (referees cross-commenting), we provide new high-resolution images with labelled membranes of both tendon cells and myoblasts (fig 2 O-P). As shown here, myoblast membranes are very closed to each other, and nuclei occupy an important part of myoblast volumes. So, we found more accurate to use the myoblast nucleus (stained with Twist antibody) to detect individual myoblasts using Imaris Spot function rather than myoblast membranes. We also believe that the distances between the center of myoblast nuclei and the tendon surface are representative of the distance between these two cell types as nucleus myoblast occupies most of the cell volume. We addressed this point in the new Suppl. Fig.5.

**Regarding the distance of 4 µm**: As mentioned in the original text, the 4 µm distance represents the average distance between myoblasts and the tendon surface in wild type discs. We do not perceive this distance as indicative of a threshold distinguishing myoblasts that interact physically with tendons from those that do not. We rather use this mean distance to quantify the distribution of myoblasts around the tendon and their dispersion/mis-distribution in Ama and Nrt knockdown leg discs. To clarify this point, we have modified the corresponding paragraph: "This result indicates that AmaKD leads to myoblasts mis-distribution around the tilt, suggesting that the reduction of Ama level could affect myoblast-tendon adhesion".

For a better understanding of how the mean distance was measured, we added a new Supplementary Figure 5 (rather than a zoom-in in the main figure as suggested by this reviewer), with a corresponding description of how this distance was measured in addition to the explanations in the material and method section.

*- Is the tendon elongation phenotype seen after Ama RNAi in muscle and in Nrt mutants due to the fact that myoblasts are further away from tendons or is it an Ama/Nrt role that is autonomous to tendons? This could be tested by assaying tendon elongation after tendon-specific Ama knock-down as shown in Figure S2.*

**Reply:** As asked by this reviewer, we have performed this experiment using Sr-Gal4 driver to induce tendon-specific Ama knockdown and assessed tendon elongation using R79D08-lexA>lexAop-GFP marker. Overall statistical analysis is now included in Fig 5F and G graphs. This analysis shows that tendon-specific Ama knockdown does not affect tendon elongation. This is in concordance with the fact that Ama knockdown in myoblasts leads to tendon defects similar to that of Nrt loss of function in tendon clearly indicating that the observed phenotypes are due to Ama's role in myoblasts. This does not exclude an additional subsequent Ama function in growing tendon precursors in later development.

**Minor:**

*- Is Figure 2Q a zoom-in from Figure 2P? If yes, it would be helpful to indicate the rough position of it in the lower magnification image.*

Figure 2Q was not a zoom-in from Figure 2P in the previous version of the paper. As stated above Fig. 2 has now been modified.

*- page 6 - w[1118] is with small "w".* modifications have been made accordingly in main and figure texts.

***REVIEWER 4:***

*The manuscript is well-organized, with clear descriptions of methods and results. The use of transcriptomic datasets and gene expression analyses provides insights into the molecular mechanisms underlying the interaction between muscle and tendon precursors.*

*The immunostaining and in situ hybridization experiments well illustrate the expression patterns of Ama and Nrt in muscle and tendon cells during leg disc development in Drosophila.*

*The functional analyses, including knockdown experiments, support the conclusion that Ama plays a crucial role in maintaining the pool of leg muscle precursor cells and coordinating tendon and muscle precursor growth.*

*The manuscript significantly enhances our understanding of cell-cell interactions in the musculoskeletal system of Drosophila. The findings have broader implications for the field of developmental biology. In general, this manuscript provides valuable insights into the molecular processes governing leg muscle and tendon development.*

We are indebted to Reviewer 4 for highlighting that our manuscript is well-organized and well-illustrated. We are also grateful to Reviewer 4 for highlighting the valuable insights of our work.

### Full Revision

***Point-by-point reply :***

- some aspects in the manuscript, for example how Ama regulates myoblast number and its interaction with the FGFR pathway, could be further explored or clarified.

**Reply:** Regarding Ama's contribution for maintaining the myoblast pool through its interaction with the FGF pathway, we demonstrate here that, contrary to what has been proposed for glial cells, Ama acts downstream of this pathway, although we emphasize that there is a synergistic effect with the MAPK pathway inhibitor, Sprouty. These findings thus reveal complex and variable regulatory mechanisms between Ama and the FGF pathway that would require specific investigation, the entirety of which appears challenging to integrate into this same publication.

- the organization of the abstract could be improved to provide a clearer and more comprehensive overview of the study. The abstract currently lacks a structured presentation of essential components such as methods, results, and conclusions. It would greatly benefit from a more systematic arrangement.

**Reply:** We have made modifications to propose a more structured abstract.
